# Acute Inhibition of Adipose Triglyceride Lipase by NG497 Dysregulates Insulin and Glucagon Secretion from Human Islets

**DOI:** 10.1101/2024.11.11.623085

**Authors:** Lucy B Kim, Siming Liu, Syreine Richtsmeier, Michał Górniak, Anamika Vikram, Yumi Imai

## Abstract

Adipose triglyceride lipase (ATGL), which catalyzes the breakdown of triglycerides in lipid droplets (LDs), plays a critical role in releasing fatty acids to support insulin secretion in pancreatic beta cells. Based on genetic downregulation of ATGL in beta cells, multiple mechanisms are proposed that acutely or chronically regulate insulin secretion. Currently, the contribution of acute versus chronic mechanisms in the regulation of insulin secretion is unclear. Also, little is known whether ATGL affects alpha cell function. Using the human-specific ATGL inhibitor, NG497, this study investigates the impact of acute inhibition of ATGL on hormone secretion from human islets. When lipolysis by ATGL was assessed via morphological differences in LDs in confocal images of beta and alpha cells, beta cells exposed to NG497 showed notable increases in LD size and number under glucose-sufficient culture. The effect of NG497 on LD accumulation in alpha cells was more prominent under fasting-simulated conditions than glucose-sufficient conditions, pointing toward a critical role for ATGL lipolysis under conditions that stimulate hormone secretion in beta and alpha cells. When exposed to NG497 acutely, human islets reduced glucose-stimulated insulin secretion mildly, particularly first-phase insulin secretion, to an extent less pronounced than the impacts of chronic ATGL downregulation. Thus, chronic mechanisms play a predominant role in reducing insulin secretion when ATGL is downregulated. Acute exposure of human islets to NG497 significantly reduced glucagon secretion at low glucose concentration, highlighting an important potential role of ATGL lipolysis in promoting hormone secretion acutely from alpha cells.

---

Adipose triglyceride lipase (ATGL, encoded by PNPLA2 gene) is a neutral lipase whose major substrates are triglycerides (TG) in lipid droplets (LDs) (1). In pancreatic beta cells, lipolysis mediated by ATGL is stimulated by glucose and supports insulin secretion through multiple mechanisms revealed by studies of beta cells after downregulation of ATGL expression (2,3). Fatty acids (FA) released from LDs can activate the GPR40 cell surface receptor and potentiate glucose-stimulated insulin-secretion (GSIS) (2). 1-monoacylglyceride (1-MAG), another product of lipolysis by ATGL, stimulates exocytosis of insulin granules through MUNC13-1 that promotes SNARE complex assembly (4). In addition, a more chronic contribution of ATGL activity for GSIS is proposed. ATGL knockout in mouse beta cells impairs mitochondrial function by reducing activity of peroxisome proliferator-activated receptors δ (PPARδ) (5). The impairment of insulin secretion after ATGL downregulation was associated with decreased syntaxin1a (STX1a) protein levels in both human pseudoislets and INS-1 cells, which is attributed to accelerated degradation of STX1a secondary to reduced palmitoylation (6). In contrast to well-established impairment of insulin secretion after genetic downregulation of ATGL in beta cell models, it is unclear the extent by which acute suppression of lipolysis by ATGL impairs insulin secretion. Acute reduction of GSIS was originally reported in rat islets treated with a pan lipase inhibitor Orlistat (7), but a subsequent report has not been consistent regarding the impact of Orlistat on GSIS (8). Moreover, Orlistat is not a specific inhibitor of ATGL creating a gap in knowledge regarding acute contribution of ATGL mediated lipolysis on GSIS in human islets (9). Another gap in knowledge is a role of ATGL in alpha cells that are proposed to utilize FA for glucagon secretion (10).

NG497 is the first human specific inhibitor of ATGL that acutely blocks lipolysis in human adipocytes and HepG2 cells at 40 nM and allows us to assess a role of ATGL through pharmacological inhibition (11). Here, we tested ATGL activity under different nutrient conditions in human beta cells and alpha cells using the change in LD morphometry as an index. In addition, the acute effect of NG497 on insulin and glucagon secretion was tested in human islets to determine the extent by which lipolysis acutely impacts human beta and alpha cell function.

## MATERIALS and METHODS

### Human islets

Institutional Review Board at University of Iowa deemed human islet experiments are not a human study. Human islets from Integrated Islet Distribution Program (IIDP), or Alberta Diabetes Institutes with reported viability and purity above 80% were cultured overnight at 37 ºC and 5% CO_2_ upon arrival for recovery from shipping. Perifusion of intact human islets were performed the next day after shipment. For morphometry of LDs, human islets dispersed by Accutase (SCR005, Millipore Sigma, St Louis, MO) were plated at approximately 35,000 cells/cm^2^ on a confocal dish coated with 50 μg/mL of Col IV (C5533 from Sigma) as reported (12), and cultured in neuronal medium (11 mM glucose) described by Phelps et al. (13) for three days. Human pseudoislets treated with lentivirus expressing shRNA against ATGL and scramble control were previously validated and reported (6).

### Morphometry of LDs

Human islets were incubated in neuronal medium containing 3 μM Bodipy 558/568 C12 (Bodipy C12, ThermoFisher, Waltham, MA) with vehicle (DMSO) or 40 μM NG497 (Focus Biomolecules, Plymouth Meeting, PA) at 37 °C at 5% CO_2_ for the last 16 hours of culture. After fixation, cells were immunostained with insulin (INS, rabbit anti-INS antibody, C27C9, Cell signaling at 1:300) or glucagon (GCG, by mouse anti-GCG antibody, K79bB10 from Sigma at 1:600) to visualize beta and alpha cells, respectively. Images of the cells were captured by a Zeiss 980 microscope to acquire z-stack images via a 63x oil lens at an interval of 0.19 μm. A set of 5 consecutive slices of the image with the largest footprint of cells were selected from each stack. The size and number of LDs were measured as published (12). The area of beta cells and alpha cells were also obtained using INS and GCG channels applying the method used for the measurement of LDs area above.

### Perifusion of Islets

BioRep Perifusion System (BioRep Technologies, Miami Lakes, FL) was used to perifuse human pseudoislets as published (14). For the measurement of insulin secretion, islets were perifused by Krebs-Ringer bicarbonate buffer (KRB) containing 2.8 mM glucose for 48 minutes followed by KRB containing 16.7 mM glucose or 30 mM KCl plus 2.8 mM glucose for indicated time. For the measurement of glucagon secretion, islets were perifused by KRB containing 3.3 mM glucose for 48 min followed by 1 mM glucose plus amino acid mixture (2 mM each of glutamine, alanine, and arginine as published in (15)) for 28 min and 7 mM glucose plus amino acid mixture for 14 min. Either vehicle (DMSO) or 40 μM NG497 was added from time 0 for all perifusion. Total insulin and glucagon contents were obtained from islets incubated overnight at 4 °C in RIPA buffer (R0278, Sigma) containing protease inhibitors. Insulin was measured using STELLUX Chemiluminescent Human Insulin ELISA (ALPCO, Macedon NY). Glucagon was measured using glucagon ELISA kit from Crystal Chem (Elk Grove Village, IL).

### Statistical analysis

Data are presented as mean ± standard error of mean (SEM) unless otherwise stated in the figure legends. Differences of numeric parameters between two groups were assessed with Student’s t-tests using Prism 10 (GraphPad, La Jolla, CA). Multiple group comparisons used One-way ANOVA with post hoc as indicated. A p < 0.05 was considered significant.

## RESULTS

### Human beta and alpha cells accumulate LDs avidly after ATGL inhibition at the conditions that stimulate their hormone secretion

Balance between the formation and degradation of LDs determines the size and number of LDs in each cell. Average size and number of LDs varied among donors, but they were overall in a similar range for beta and alpha cells cultured in 11 mM glucose medium (Fig. 1A-C). We assessed the proportion of LDs being degraded by ATGL under 11 mM glucose medium by measuring the increase in size and number of LDs after NG497 treatment in beta and alpha cells. For both beta and alpha cells, the inhibition of lipolysis overnight by NG497 caused statistically significant increases in the size and number of LDs contributing to the increase in total area occupied by LDs corrected for cell area (Fig. 1A-D). For beta cells, the prominent accumulation of LDs by NG497 agrees with the increase of LDs observed in human beta cells when ATGL is downregulated by shRNA and is in line with previous reports of ATGL activation by glucose in beta cells (6,16). However, the impact of ATGL inhibition on LD accumulation at 11 mM glucose was less prominent in alpha cells, shown as lower fold change of LD size, LD number, and total area occupied by LDs compared with beta cells (Fig. 1B-D, right panels). Smaller LD accumulation by NG497 implicates that ATGL activity is lower in alpha cells than beta cells under glucose sufficient conditions.

**Figure 1.**
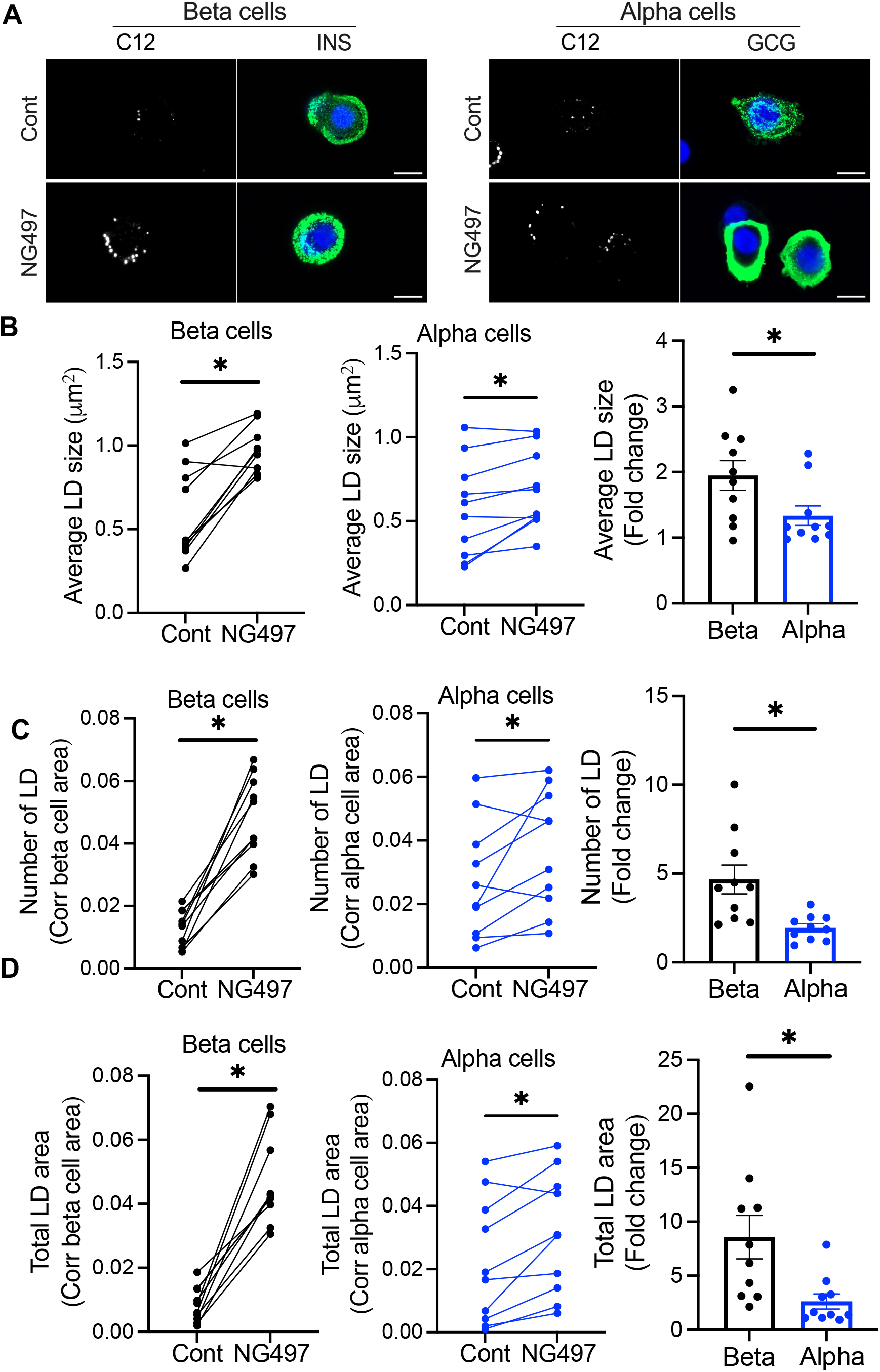
NG497 promotes LD accumulation more prominently in human beta cells than alpha cells under 11 mM glucose. A) Representative confocal images of human beta and alpha cells cultured in 11 mM glucose medium with overnight addition of vehicle (Cont) or NG497. Cells were stained with DAPI (blue), BODIPY 558/568 C12 (C12, white) to visualize LDs, and INS or GCG (green) to determine beta and alpha cell identity. Scale bar, 10 μm. B) Average size of LDs, C) average number of LDs corrected for insulin- or glucagon-positive area, and D) average total LD area corrected for insulin- or glucagon-positive area in Cont and NG497-treated beta and alpha cells. Left and middle panels: Each dot represents one donor and values from the same donor are connected by a line. Right panel: Fold change of NG497-treated cells over Cont. Data are mean ± SEM. n = 10 donors *; p<0.05 by Student’s t-test.

Considering that glucagon secretion from alpha cells is more active during fasting when glucose level is low and FA availability is increased (10), we tested whether the disruption of lipolysis affects the degree of LD accumulation more prominently in a culture condition similar to fasting. Human islets with and without NG497 treatment were cultured under 11 mM glucose or 2.8 mM glucose plus 0.13 mM oleic acid and 0.27 mM palmitic acid (LoG +FA) loading overnight before fixation (Fig. 2A). For three independent donor islets tested, NG497 increased average LD size to a similar extent both in 11 mM glucose and LoG + FA conditions (Fig. 2B). However, NG497 increased the number of LDs and total LD area per cell area more prominently under LoG + FA conditions than under 11 mM glucose (Fig. 2C-D). Thus, LD degradation by ATGL lipolysis in alpha cells is more active under LoG + FA conditions compared with 11 mM glucose.

**Figure 2.**
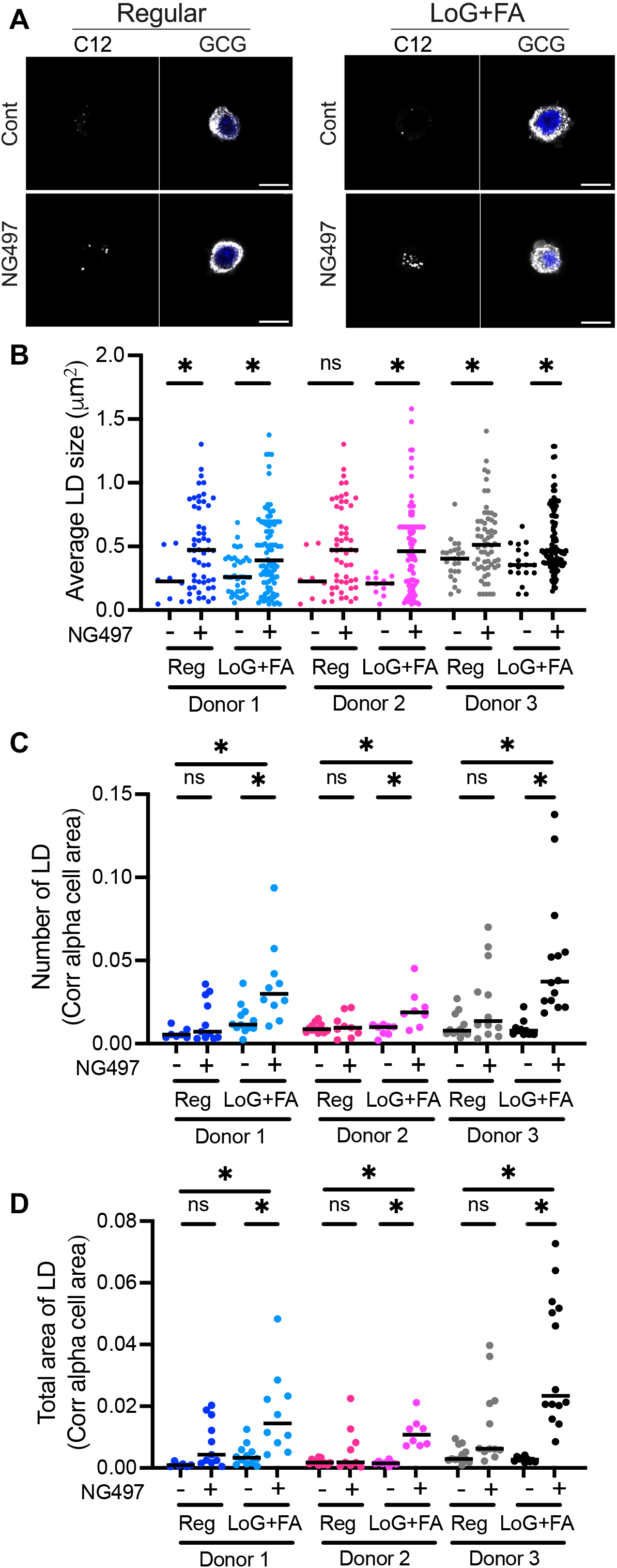
Accumulation of LDs in human alpha cells increases under low glucose and fatty acid loading. A) Representative confocal images of alpha cells cultured in medium containing 11 mM glucose (Reg) or 2.8 mM glucose with 0.13 mM oleic acid plus 0.27 mM palmitic acid loading (LoG + FA) with addition of vehicle (Cont) or NG497 for overnight. Alpha cells were identified via DAPI (blue) and GCG-staining (white) while LDs were visualized with BODIPY 558/568 C12 (C12, white). Scale bar, 10 μm. B) Size distribution of LDs in alpha cells, C) the number of LDs corrected for glucagon-positive area, and D) the total LD area occupied per glucagon-positive area for three donor islets. Each dot represents an individual LD for B) and an individual image for C) and D). n=6 to 14 images analyzed. Median is indicated by line. *; p-value<0.05 by One-way ANOVA with Sidak’s multiple comparison test.

### Acute ATGL inhibition predominantly impairs first phase of glucose-stimulated insulin secretion from human beta cells

To compare the impact of acute vs prolonged suppression of ATGL activity on GSIS, GSIS was measured in human pseudoislets in which ATGL was downregulated by shRNA and control pseudoislets acutely exposed to NG497. qPCR confirmed downregulation of ATGL to 27.1 ± 6.3% of scramble control in shATGL treated pseudoislets (n=4, p<0.05 by Student’s t test). We saw clear reduction of GSIS in shATGL treated pseudoislets as we reported before (Fig. 3A, B) (6). NG497 caused reduced GSIS in some donors but did not reach statistical significance after 4 donor pseudoislets were tested. Insulin contents at the end of perifusion did not differ between three groups (data not shown). Perifusion in human intact islets from additional donors (total 7 donors) revealed mild reduction of GSIS by NG497 exposure with more prominent change in the first phase of insulin secretion than the second phase (Fig. 3C-E). NG497 did not change KCl stimulated insulin secretion or insulin contents at the end of perifusion (Fig. 3F, G).

**Figure 3.**
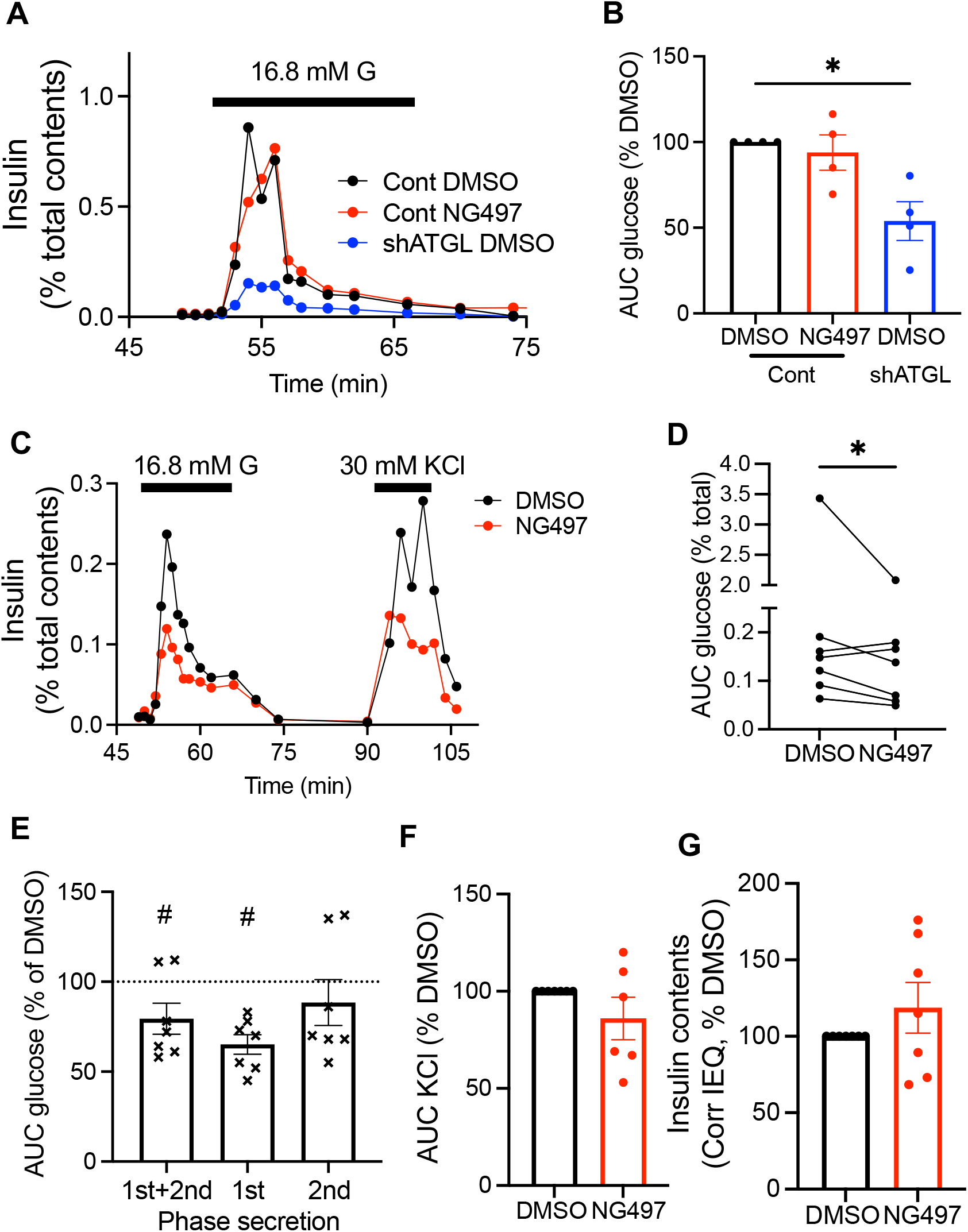
ATGL suppression dysregulates glucose-stimulated insulin secretion from human islets. A-B) Human pseudoislets transduced with lentivirus expressing shATGL (shATGL) or scramble control (Cont) were perifused with 2.8 mM glucose KRB containing 0.1% DMSO or 40 μM NG497 for 48 min followed by 16.8 mM glucose (G) for indicated time. Insulin secretion was corrected for insulin contents. A) A representative profile and B) area under the curve (AUC) of glucose response corrected for insulin contents and expressed by taking the value for Cont DMSO as 100% for each donor islets. n=4 donor. *; p<0.05 by repeated measure One-way ANOVA with Dunnette’s test. C-G) Human islets were perifused as in (A) followed by 30 mM KCl for indicated time. Insulin secretion was corrected for insulin contents. C) A representative profile. D) AUC of glucose response. Values from the same donor are connected by a line. E) First (1st) and second (2nd) phases of glucose-stimulated insulin secretion expressed taking the value for DMSO as 100% for each donor islets. F) AUC of KCl stimulated insulin secretion corrected for insulin contents and expressed by taking the value for DMSO as 100% for each donor islets. G) Insulin contents corrected for islet equivalent (IEQ) and expressed taking value for DMSO as 100% for each donor islets. C-G) n=7 donors. Data are mean ± SEM. D)* and F)#; p<0.05 vs DMSO by paired Student’s t test.

### Acute ATGL inhibition dysregulated glucagon secretion from human islets

Human islets showed marked increase in glucagon secretion in response to 1 mM glucose in the presence of amino acid mixture that lasted ∼ 10 min. Subsequent exposure to 7 mM glucose resulted in a smaller peak of glucagon secretion in control human islets (Fig. 4A). Perifusion in the presence of NG497 did not alter glucagon contents at the end of perifusion (Fig. 4B).

**Figure 4.**
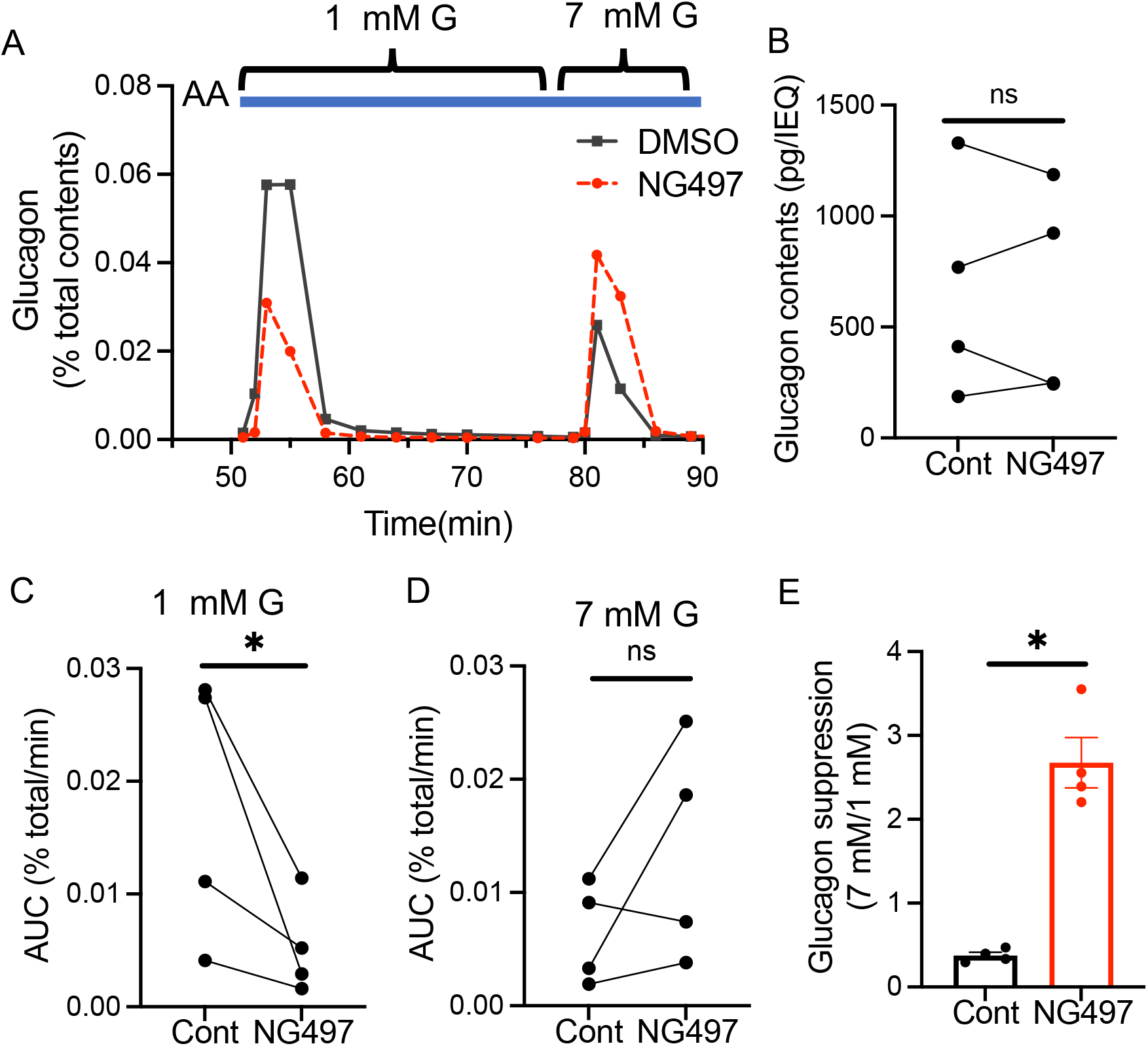
ATGL inhibition by NG497 dysregulates glucagon secretion from human islets. A-E) Human islets were first perifused by 3.3 mM glucose KRB without amino acids (AA). At indicated time, perifusion buffer was changed to KRB containing AA and 1 mM glucose (G) followed by 7 mM G. 0.1% DMSO or 40 μM NG497 was present throughout the perifusion. A) A representative profile. B) Glucagon contents per one islet equivalent (IEQ). Values from the same donor are connected by a line. C-D) Area under the curve (AUC) of glucagon peak response during C) 1 mM glucose and D) 7 mM glucose was corrected for glucagon contents and divided by the length of peak (min). E) The ratio of D) over C) as the suppression of glucagon by glucose. n=4 donors. Data are mean ± SEM. ns; not significant, *; p<0.05 by paired Student’s t test.

Notably, NG497 blunted glucagon secretion at 1 mM glucose plus amino acid mixture significantly (Fig. 4C). In contrast, glucagon secretion at 7 mM glucose plus amino acid mixture was maintained in the presence of NG497 and was higher than DMSO control in three out of four donors (Fig. 4C, D). Collectively, 7 mM glucose suppressed glucagon secretion by 64% over 1 mM glucose in control human islets but it increased 2.7-fold in NG497-exposed human islets (Fig. 4E).

## DISCUSSION

As glucose-stimulated lipolysis by ATGL is reported to be blunted in human islets from donors with type 2 diabetes, it is important to understand the mechanism by which ATGL regulates GSIS in human islets (6). Direct actions of lipolytic metabolites on the insulin secretory pathway (FA on GPR40, 1-MAG on MUNC13-1), transcriptional activation through PPARδ, and protein stabilization by palmitoylation (STX1a) are potential mechanisms proposed based on genetic downregulation of ATGL in beta cell models (4-6). In the present study, we aimed to gauge the impact of acute versus chronic ATGL inhibition on GSIS in human beta cells using a human-specific ATGL inhibitor, NG497 (11). NG497 inhibits GSIS mildly, primarily during the first phase in human islets, an extent that is smaller than one elicited by shRNA mediated downregulation of ATGL in human pseudoislets (Fig. 3)(6). Thus, chronic targets such as PPARδ and protein palmitoylation may predominantly contribute to impaired GSIS after ATGL downregulation in beta cells. This is in agreement with statistically significant but limited recovery of GSIS by the supplementation of 1-MAG in ATGL downregulated human pseudoislets (6).

Glucagon secretion is reported to be dysregulated early during the development of type 2 diabetes (17). The current study demonstrated functional significance of ATGL in glucagon secretion from human islets. LDs accumulated more prominently after ATGL inhibition in beta cells than alpha cells under glucose-sufficient culture, indicating that ATGL activity is lower in alpha cells at this condition. However, ATGL avidly contributes to LD degradation in alpha cells in the condition simulating fasting (LoG + FA). In vivo, glucagon secretion increases during hypoglycemia as a counter regulatory mechanism to increase hepatic gluconeogenesis and after dietary protein ingestion to regulate circulating amino acid levels (18-20). Compared with glucose-centered regulation of insulin secretion, the regulation of glucagon secretion by nutrients is multifactorial and has not reached consensus (10). FA are reported to increase glucagon secretion by raising cytosolic calcium in rat and mouse islets (21,22). More recently, it is proposed that glucose levels alter glucagon secretion by switching energy production between the TCA cycle fueled by glucose and beta oxidation of FA, for which CPT1a is a key node of regulation as a transporter for FA into mitochondria (23,24). While external FA can be utilized directly for FA oxidation to alter glucagon secretion, FA can also traffic through LDs via incorporation into TG and then subsequent release by ATGL (1). Thus, the reduction of glucagon secretion at 1 mM glucose by NG497 may be due to decreased FA availability for FA oxidation. Although there was not statistical significance, NG497 showed a trending increase in glucagon secretion at 7 mM glucose (Fig. 4D). It is proposed that palmitoylation of the K_ATP_ channel by FA maintains the closed status of the channel to reduce glucagon secretion (25). It is plausible that reduced lipolysis may increase glucagon secretion by reducing palmitoylation of the K_ATP_ channel. Further studies are required to address the mechanisms by which ATGL inhibition dysregulates glucagon secretion, including testing the impact of NG497 on FA oxidation or the K_ATP_ channel.

The current study has some limitations. We used NG497 as a human specific acute inhibitor of ATGL (11). LD morphometry of human beta cells exposed to NG497 overnight showed the increase in size and number of LDs indicative of impaired ATGL function supporting its activity against ATGL (6). Significant change we observed in glucagon secretion supports that NG497 elicited its action after 48 min of preincubation. However, we did not measure the activity of ATGL under NG497 treatment directly to establish that NG497 effectively blocked ATGL during the time frame of perifusion. Although glycerol release is often used to measure lipolysis, glycerol is not reliable measurement of lipolysis in beta cells that are known to secrete glycerol metabolized through glycerol-3 phosphate phosphatase, bypassing lipolysis (26). Also, it is difficult to measure lipolysis in beta cells and alpha cells separately in human islets. While human beta cells and alpha cells could be sorted from dispersed human islets, the significant loss in cells would make it difficult to measure lipolysis directly. However, future studies are needed to confirm that dysregulation of glucagon secretion by NG497 is elicited by the specific inhibition of ATGL.

In summary, the current study indicates that ATGL mediated lipolysis is an active process in both human beta and alpha cells and plays a critical role for the regulation of both insulin and glucagon secretion in human islets.

## Disclosure

The authors have nothing to disclose.

## Data availability

Some data sets generated during and/or analyzed during this study are not publicly available but are available from the corresponding author on reasonable request.

## Acknowledgement

YI is supported by the National Institutes of Health (R01-DK090490), Department of Veteran Affairs (I01 BX005107), and Fraternal Orders of Eagles Diabetes Research Center at the University of Iowa. A Zeiss 980 confocal microscope located in the University of Iowa Central Microscopy Research Facility (CMRF) was funded by the Roy J Carver Charitable Trust. MG was supported by the Erasmus+ mobility exchange program (KA 131). A part of human pancreatic islets was provided by the NIDDK-funded Integrated Islet Distribution Program (IIDP) (RRID:SCR _014387) at City of Hope, NIH Grant # U24DK098085. Human islets were also provided by the Alberta Diabetes Institute Islet Core at the University of Alberta in Edmonton with the assistance of the Human Organ Procurement and Exchange (HOPE) program, Trillium Gift of Life Network (TGLN), and other Canadian organ procurement organizations.

## References

1. Grabner GF, Xie H, Schweiger M, Zechner R. Lipolysis: cellular mechanisms for lipid mobilization from fat stores. Nat Metab. 2021;3(11):1445–1465.

2. Prentki M, Corkey BE, Madiraju SRM. Lipid-associated metabolic signalling networks in pancreatic beta cell function. Diabetologia. 2020;63(1):10–20.

3. Tong X, Liu S, Stein R, Imai Y. Lipid Droplets’ Role in the Regulation of beta-Cell Function and beta-Cell Demise in Type 2 Diabetes. Endocrinology. 2022;163(3):bqac007.

4. Zhao S, Mugabo Y, Iglesias J, Xie L, Delghingaro-Augusto V, Lussier R, Peyot ML, Joly E, Taib B, Davis MA, Brown JM, Abousalham A, Gaisano H, Madiraju SR, Prentki M. alpha/beta-Hydrolase domain-6-accessible monoacylglycerol controls glucose-stimulated insulin secretion. Cell Metab. 2014;19(6):993–1007.

5. Tang T, Abbott MJ, Ahmadian M, Lopes AB, Wang Y, Sul HS. Desnutrin/ATGL activates PPARdelta to promote mitochondrial function for insulin secretion in islet beta cells. Cell Metab. 2013;18(6):883–895.

6. Liu S, Promes JA, Harata M, Mishra A, Stephens SB, Taylor EB, Burand AJ, Jr., Sivitz WI, Fink BD, Ankrum JA, Imai Y. Adipose Triglyceride Lipase Is a Key Lipase for the Mobilization of Lipid Droplets in Human beta-Cells and Critical for the Maintenance of Syntaxin 1a Levels in beta-Cells. Diabetes. 2020;69(6):1178–1192.

7. Mulder H, Yang S, Winzell MS, Holm C, Ahren B. Inhibition of lipase activity and lipolysis in rat islets reduces insulin secretion. Diabetes. 2004;53(1):122–128.

8. Cantley J, Burchfield JG, Pearson GL, Schmitz-Peiffer C, Leitges M, Biden TJ. Deletion of PKCepsilon selectively enhances the amplifying pathways of glucose-stimulated insulin secretion via increased lipolysis in mouse beta-cells. Diabetes. 2009;58(8):1826–1834.

9. Iglesias J, Lamontagne J, Erb H, Gezzar S, Zhao S, Joly E, Truong VL, Skorey K, Crane S, Madiraju SR, Prentki M. Simplified assays of lipolysis enzymes for drug discovery and specificity assessment of known inhibitors. J Lipid Res. 2016;57(1):131–141.

10. Armour SL, Stanley JE, Cantley J, Dean ED, Knudsen JG. Metabolic regulation of glucagon secretion. J Endocrinol. 2023;259(1):e230081.

11. Grabner GF, Guttenberger N, Mayer N, Migglautsch-Sulzer AK, Lembacher-Fadum C, Fawzy N, Bulfon D, Hofer P, Zullig T, Hartig L, Kulminskaya N, Chalhoub G, Schratter M, Radner FPW, Preiss-Landl K, Masser S, Lass A, Zechner R, Gruber K, Oberer M, Breinbauer R, Zimmermann R. Small-Molecule Inhibitors Targeting Lipolysis in Human Adipocytes. J Am Chem Soc. 2022;144(14):6237–6250.

12. Brennecke BR, Yang U, Liu S, Ilerisoy FS, Ilerisoy BN, Joglekar A, Kim LB, Peachee SJ, Richtsmeier SL, Stephens SB, Sander EA, Strack S, Moninger TO, Ankrum JA, Imai Y. Utilization of commercial collagens for preparing well-differentiated human beta cells for confocal microscopy. Front Endocrinol (Lausanne). 2023;14:1187216.

13. Phelps EA, Cianciaruso C, Santo-Domingo J, Pasquier M, Galliverti G, Piemonti L, Berishvili E, Burri O, Wiederkehr A, Hubbell JA, Baekkeskov S. Advances in pancreatic islet monolayer culture on glass surfaces enable super-resolution microscopy and insights into beta cell ciliogenesis and proliferation. Sci Rep. 2017;7:45961.

14. Harata M, Liu S, Promes JA, Burand AJ, Ankrum JA, Imai Y. Delivery of shRNA via lentivirus in human pseudoislets provides a model to test dynamic regulation of insulin secretion and gene function in human islets. Physiol Rep. 2018;6(20):e13907.

15. Singh B, Khattab F, Gilon P. Glucose inhibits glucagon secretion by decreasing [Ca(2+)](c) and by reducing the efficacy of Ca(2+) on exocytosis via somatostatin-dependent and independent mechanisms. Mol Metab. 2022;61:101495.

16. Prentki M, Madiraju SR. Glycerolipid/free fatty acid cycle and islet beta-cell function in health, obesity and diabetes. Molecular and cellular endocrinology. 2012;353(1-2):88-100.

17. Kohlenberg JD, Laurenti MC, Egan AM, Wismayer DS, Bailey KR, Cobelli C, Man CD, Vella A. Differential contribution of alpha and beta cell dysfunction to impaired fasting glucose and impaired glucose tolerance. Diabetologia. 2023;66(1):201–212.

18. Capozzi ME, D’Alessio DA, Campbell JE. The past, present, and future physiology and pharmacology of glucagon. Cell Metab. 2022;34(11):1654–1674.

19. Hahn HJ, Ziegler M. Investigations on isolated islets of langerhans in vitro. 16.Modification of the glucose-dependent inhibition of glucagon secretion. Biochim Biophys Acta. 1977;499(3):362–372.

20. Liu L, Dattaroy D, Simpson KF, Barella LF, Cui Y, Xiong Y, Jin J, Konig GM, Kostenis E, Roman JC, Kaestner KH, Doliba NM, Wess J. Gq signaling in alpha cells is critical for maintaining euglycemia. JCI Insight. 2021;6(24).

21. Olofsson CS, Salehi A, Gopel SO, Holm C, Rorsman P. Palmitate stimulation of glucagon secretion in mouse pancreatic alpha-cells results from activation of L-type calcium channels and elevation of cytoplasmic calcium. Diabetes. 2004;53(11):2836–2843.

22. Fujiwara K, Maekawa F, Dezaki K, Nakata M, Yashiro T, Yada T. Oleic acid glucose-independently stimulates glucagon secretion by increasing cytoplasmic Ca2+ via endoplasmic reticulum Ca2+ release and Ca2+ influx in the rat islet alpha-cells. Endocrinology. 2007;148(5):2496–2504.

23. Briant LJB, Dodd MS, Chibalina MV, Rorsman NJG, Johnson PRV, Carmeliet P, Rorsman P, Knudsen JG. CPT1a-Dependent Long-Chain Fatty Acid Oxidation Contributes to Maintaining Glucagon Secretion from Pancreatic Islets. Cell Rep. 2018;23(11):3300–3311.

24. Armour SL, Frueh A, Chibalina MV, Dou H, Argemi-Muntadas L, Hamilton A, Katzilieris-Petras G, Carmeliet P, Davies B, Moritz T, Eliasson L, Rorsman P, Knudsen JG. Glucose Controls Glucagon Secretion by Regulating Fatty Acid Oxidation in Pancreatic alpha-Cells. Diabetes. 2023;72(10):1446–1459.

25. Veprik A, Denwood G, Liu D, Bany Bakar R, Morfin V, McHugh K, Tebeka NN, Vetterli L, Yonova-Doing E, Gribble F, Reimann F, Hoehn KL, Hemsley PA, Ahnfelt-Ronne J, Rorsman P, Zhang Q, de Wet H, Cantley J. Acetyl-CoA-carboxylase 1 (ACC1) plays a critical role in glucagon secretion. Commun Biol. 2022;5(1):238.

26. Attane C, Peyot ML, Lussier R, Poursharifi P, Zhao S, Zhang D, Morin J, Pineda M, Wang S, Dumortier O, Ruderman NB, Mitchell GA, Simons B, Madiraju SR, Joly E, Prentki M. A beta cell ATGL-lipolysis/adipose tissue axis controls energy homeostasis and body weight via insulin secretion in mice. Diabetologia. 2016;59(12):2654–2663.

